# A Chemical-mechanical Coupled Model Predicts Roles of Spatial Distribution of Morphogen in Maintaining Tissue Growth

**DOI:** 10.1101/2022.06.28.497907

**Authors:** Alireza Ramezani, Samuel Britton, Roya Zandi, Mark Alber, Ali Netmatbakhsh, Weitao Chen

## Abstract

The exact mechanism controlling cell growth remains a grand challenge in developmental biology and regenerative medicine. The *Drosophila* wing disc tissue serves as an ideal biological model to study growth regulation due to similar features observed in other developmental systems. The mechanism of growth regulation in the wing disc remains a subject of intense debate. Most existing models to study tissue growth focus on either chemical signals or mechanical forces only. Here we developed a multiscale chemical-mechanical coupled model to test a growth regulation mechanism depending on the spatial range of the morphogen gradient. By comparing the spatial distribution of cell division and the overall shape of tissue obtained in the coupled model with experimental data, our results show that the distribution of the Dpp morphogen can be critical in resulting tissue size and shape. A larger tissue size with a faster growth rate and more symmetric shape can be achieved if the Dpp gradient spreads in a larger domain. Together with the absorbing boundary conditions, the feedback regulation that downregulates Dpp receptors on the cell membrane allows the further spread of the morphogen away from its source region, resulting in prolonged tissue growth at a more spatially homogeneous growth rate.

**Summary Statement:** A multiscale chemical-mechanical model was developed by coupling submodels representing dynamics of a morphogen gradient at the tissue level, intracellular chemical signals, and mechanical properties at the subcellular level. By applying this model to study the *Drosophila* wing disc, it was found that the spatial range of the morphogen gradient affected tissue growth in terms of the growth rate and the overall shape.

## Introduction

Understanding mechanisms underlying proper tissue growth and shape formation in an embryo are among the most important unanswered questions in developmental biology and remain elusive for various biological systems. The growth of tissues and organs always exhibits the property of self-organization, which means cell proliferation is under precise control individually to give rise to a robust tissue size and specific shape as integrity. This process is often independent of cell size, length of the cell cycle, and robust to external perturbations as observed in wound healing and tissue regeneration (Boulan et al., 2015; Bryant and Levinson, 1985; Harrison, 1924; Vollmer et al., 2017; Twitty and Schwind, 1931; Zhu et al., 2020). Uncontrolled cell growth often leads to abnormal development or fatal diseases such as cancer.

During tissue development, chemical cues are found to be critical to the regulation of cell proliferation and tissue shape formation. A variety of molecules, from extracellular ligands to intracellular proteins, have been identified as growth regulators in different biological systems. For example, transforming growth factor Beta (TGF-β), a member of the growth factor superfamily, has been found to regulate the growth in multiple animal organs (Heldin et al., 1997; Siegel and Massagué, 2003). In particular, bone morphogenic proteins (BMP) are members of the TGF-β family and play essential roles in establishing the basic embryonic body plan for the tissue development in vertebrates (Bellusci et al., 1996; Hammerschmidt et al., 1996; Jones et al., 1991; Lyons et al., 1990; Martínez-Barberá et al., 1997). Disruption of BMP signals can affect the growth rate and pattern formation, leading to disorders in adult tissues (Ide et al., 1997; Zhao, 2003). On the other hand, in addition to the central core of the growth control machinery which depends on chemical cues, cell mechanics play a fundamental role in shaping a tissue (Chien et al., 1998; Huang and Ingber, 1999; Huiskes et al., 2000; Jain et al., 2014; Kitterman, 1996; Mao et al., 2013; Minc et al., 2009; Shraiman, 2005). Each cell has a complex mechanical architecture that not only shapes itself as integrity but also allows it to sense the physical surroundings. For example, cytoskeletal tension in one cell can be affected by differential growth associated with neighboring cells and modulate intracellular molecular signals to regulate growth as feedback (Pan et al., 2016). Cell deformation can be induced by mechanical forces such as the adhesion to the extracellular matrix (ECM), contractility in the cytoskeleton, and the cell-cell adhesion, leading to physical changes of nuclei and an alteration in gene expression to switch cell fate between growth, differentiation, and apoptosis (Werfel et al., 2013). Therefore, it is necessary to consider both chemical and mechanical cues, as well as the interplay between them, to study tissue development.

*Drosophila* wing disc, a primordial epithelial organ that later becomes the adult wing, as shown in Fig. 1A, serves as a classic model to study tissue growth regulation, due to its simple geometry, a limited number of cells, and fast growth. Additionally, the well-established molecular signaling network involved in this system contains multiple conserved molecules critical to other developing systems in mammals (Siegel and Massagué, 2003). Understanding the mechanism of growth regulation in *Drosophila* wing disc is substantial toward understanding limb development in mammals. In this tissue, Decapentaplegic (Dpp), a homolog of BMP, forms a spatial gradient across the anterior-posterior (AP) axis of the tissue to establish and maintain domains of multiple target genes that specify different compartments in the adult tissue (Fig. 1B,C). For individual cells, a signal transduction cascade converts local concentrations of Dpp into an intracellular phosphorylated Mad (pMad) concentration through binding with receptors on the membrane. pMAD protein is also commonly observed in other systems and correlated to several cancers in humans (Heldin et al., 1997). Based on the level of pMad, different genes are activated along the AP axis of the imaginal wing disc to establish the pattern and regulate growth. In terms of mechanical properties, a wing disc consists of a flat sheet of cells with E-cadherin responsible for cell-cell adhesion between neighboring cells. Inside individual cells, actomyosin is dynamically rearranged to give rise to different levels of contractility, which links to multiple cellular functions including nuclear motion and vesicle trafficking (Masedunskas et al., 2011; Meitinger and Palani, 2016). Moreover, actin networks in the cytoplasm, as a major component of the cytoskeleton, provide structural support to each cell and determine cell shapes together with the cytoplasm. More recently, it has been observed that chemical signals can affect cell mechanics directly by controlling the subcellular distribution of the small GTPase Rho1 and the regulatory light chain of non-muscle myosin (Widmann and Dahmann, 2009). Dpp signal promotes the compartmentalization of Rho1 and myosin, which leads to the contraction of actomyosin filaments and an increase in cortical tension. This suggests the interaction between chemical signals and mechanical properties plays an important role in shaping cells and hence the overall wing disc tissue.

**Figure 1.**
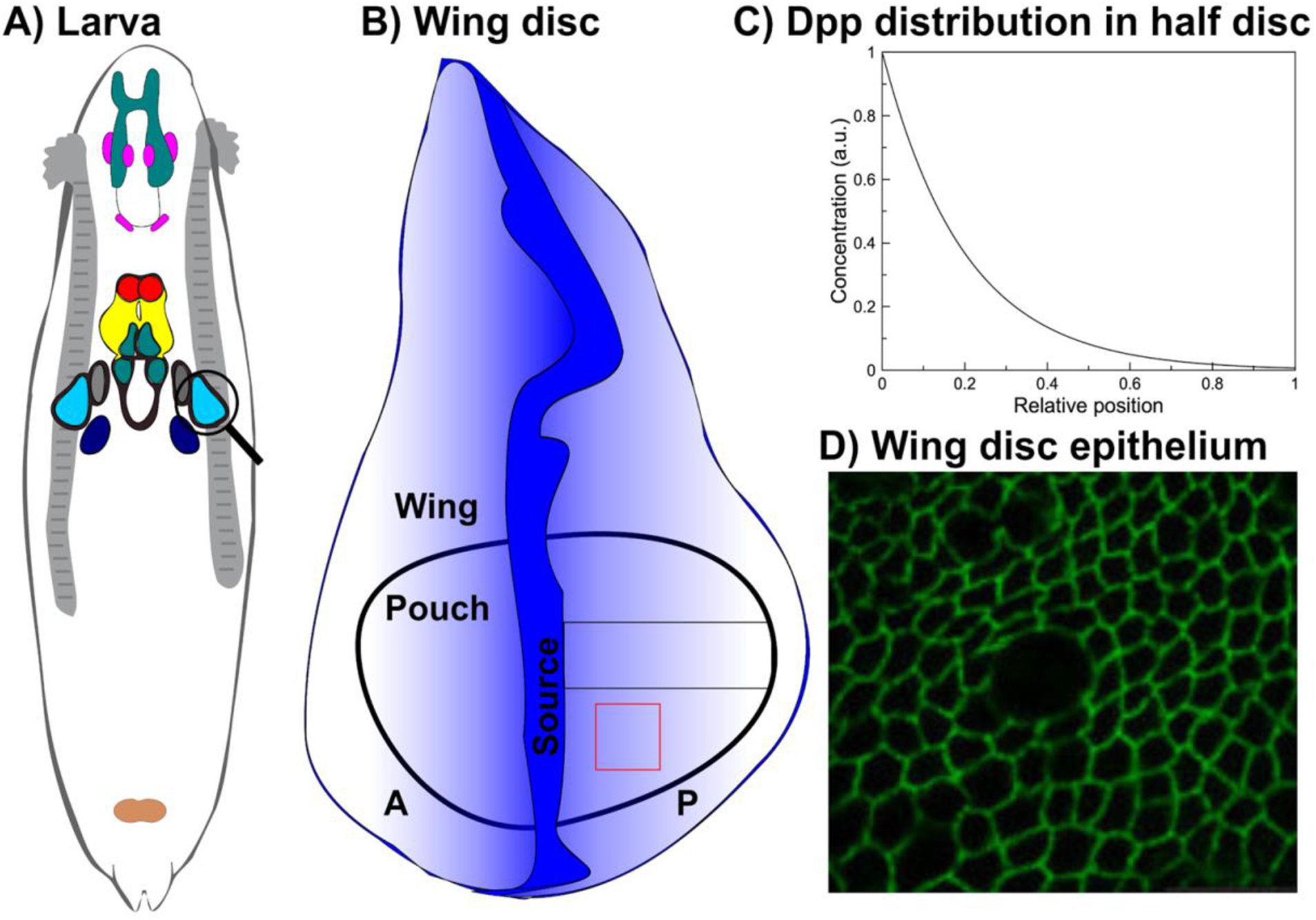
A) Illustrative picture of Drosophila larva with wing disc tissue circled. B) Zoom-in view of the Drosophila imaginal wing disc. The blue color denotes the Dpp morphogen gradient. C) Schematic profile of the Dpp morphogen in half wing disc. Its distribution follows an exponential shape as observed in experiments. D) Configuration of epithelial cells in the wing disc pouch. The image has been reproduced from Gibson et al., (2014).

Several hypotheses for growth regulation in the wing disc have been proposed so far, with many of them depending on chemical signal cues only. In Wartlick et al., (2011), it was suggested that cells have memory and will divide if the temporal variation in the Dpp signal reaches a certain threshold value. However, more recent experiments have shown that Dpp signal is not always required for growth, since removing Dpp from the center of the tissue at some stage during the development doesn’t affect the growth (Akiyama and Gibson, 2015). Moreover, mechanical properties have been shown to be critical in regulating growth based on measurements of cell stress in experiments (Pan et al., 2016). Although substantial data suggests that morphogens are pivotal in regulating growth, the underlying mechanism remains controversial and uncertain.

A variety of modeling approaches have been developed to study tissue growth in different biological systems, including cell lineage in epithelia (Chou et al., 2010; Du et al., 2015; Lo et al., 2009; Zhang et al., 2012) and tumor growth (Adam, 1986; Cristini et al., 2003; Wise et al., 2008). To include cell mechanics, one type of models uses discrete particles to represent individual cells, which allows one to model cell growth, cell division, and cell-cell interaction. In this type of models, each cell is usually represented by a single particle (agent-based model), multiple particles on the cell membrane (multi-agent-based model), a polygon (vertex-based model), or multiple particles on the cell membrane and in the cytoplasm (subcellular element method). In particular, multi-agent-based models and sub-cellular element models can describe biologically relevant cell shapes with greater flexibility due to multiple nodes involved. Another type of models is based on the finite element framework, coupled with continuum mechanics principles. This type of models focuses more on tissue growth without subcellular details. To include chemical signaling networks, it is common to use continuous models in which the dynamics of chemical signals are captured by a system of differential equations. This kind of approach usually involves moving boundary problems for capturing tissue growth that are challenging to solve numerically. It can be handled using a Lagrangian framework, immersed boundary method (Peskin, 2002), level set method (Osher and Sethian, 1988), or other similar approaches. Most existing models for studying tissue growth focus on either chemical signals or mechanical properties. As suggested by recent data, exploring the growth regulation in a wing disc requires a model that includes both chemical and mechanical factors, as well as the interactions between them. Several coupled chemical-mechanical models have been developed recently and gained lots of attention. In Aegerter-Wilmsen et al., (2012), both chemical signals and mechanical cues were considered, however, fixed morphogen gradients were adopted without considering the temporal dynamics or subcellular activities. Vertex-based models have been coupled at the cell level with diffusive molecules (Tanaka et al., 2015) or intracellular gene expression (Heldin et al., 1997; Siegel and Massagué, 2003) to study tissue development. The subcellular element model has also been coupled with chemical signals without distinguishing cell membrane and cytoplasm (Newman, 2005; Sandersius and Newman, 2008). Those existing models provide novel insights into the growth regulation in different systems. As far as we know, very few of them consider subcellular details and the interaction between chemical signals and mechanical forces, which is critical in regulations of individual cell behavior and tissue growth.

In this study, we developed a multiscale coupled chemical-mechanical model where the mechanical submodel described cell mechanics at the subcellular level, and the chemical signaling network described both morphogen gradient at the tissue level and the intracellular gene regulatory network at the cell level. This coupled chemical-mechanical model was then applied to study growth regulation in *Drosophila* wing imaginal disc. In particular, we incorporated a cell division rule proposed in Wartlick et al., (2011), in which cells enter the mitotic phase and divide when the Dpp concentration is increased by 50% compared with that at the beginning of the cell cycle. Following this hypothesis, the morphogen gradient with different decay lengths was tested in the model to simulate tissue growth. We found that, under the specific cell division rule, a morphogen distribution with a larger decay length could maintain the tissue growth longer to reach a more symmetric shape at a more spatially homogeneous growth rate as observed in experiments. Together with the absorbing boundary condition, feedback regulation of the downstream signal to inhibit the synthesis of cell membrane receptors facilitated tissue growth by increasing the decay length of the morphogen indirectly. This subcellular coupled chemical-mechanical model could be applied to test various hypotheses on growth regulation involved in other biological systems.

## Results

### Description of the multiscale coupled chemical-mechanical model

During tissue development, both chemical signals and mechanical forces play essential roles in regulating cell growth. We have developed a novel model to integrate both chemical and mechanical factors and the interactions between them at the subcellular level. This chemical-mechanical model employs a subcellular element particle-based method for the mechanical submodel and a system of differential equations as the model for chemical signals coupled in both space and time. Details of each submodel are provided in Materials and Methods.

#### Spatial Coupling of Mechanical and Chemical Submodels

The spatial coupling of the chemical signaling submodel and the mechanical submodel was achieved through a dynamic triangular mesh over the spatial domain of individual cells, which also covered the entire tissue. Such a dynamic mesh was constructed based on the discrete nodes representing cell membranes obtained in the mechanical submodel (Fig. 2A). Common edges and junction points shared by neighboring cells were identified as the edges and vertices of triangles, respectively (Fig. 2A’-A”). Together with cell centers, they gave rise to a triangular mesh covering individual cells (Fig. 2A’’’, B-C). More details about this mesh generator are provided in Materials and Methods. The chemical signaling submodel described in Eqs. 10–13 in general was then computed over the mesh at a frequency determined in the temporal coupling. Distributions of chemical signal concentrations were obtained at individual cells and tissue level (Fig. 2E). Meanwhile, cell averages of the chemical signals were fed into the mechanical submodel to direct cell growth or division.

**Figure 2.**
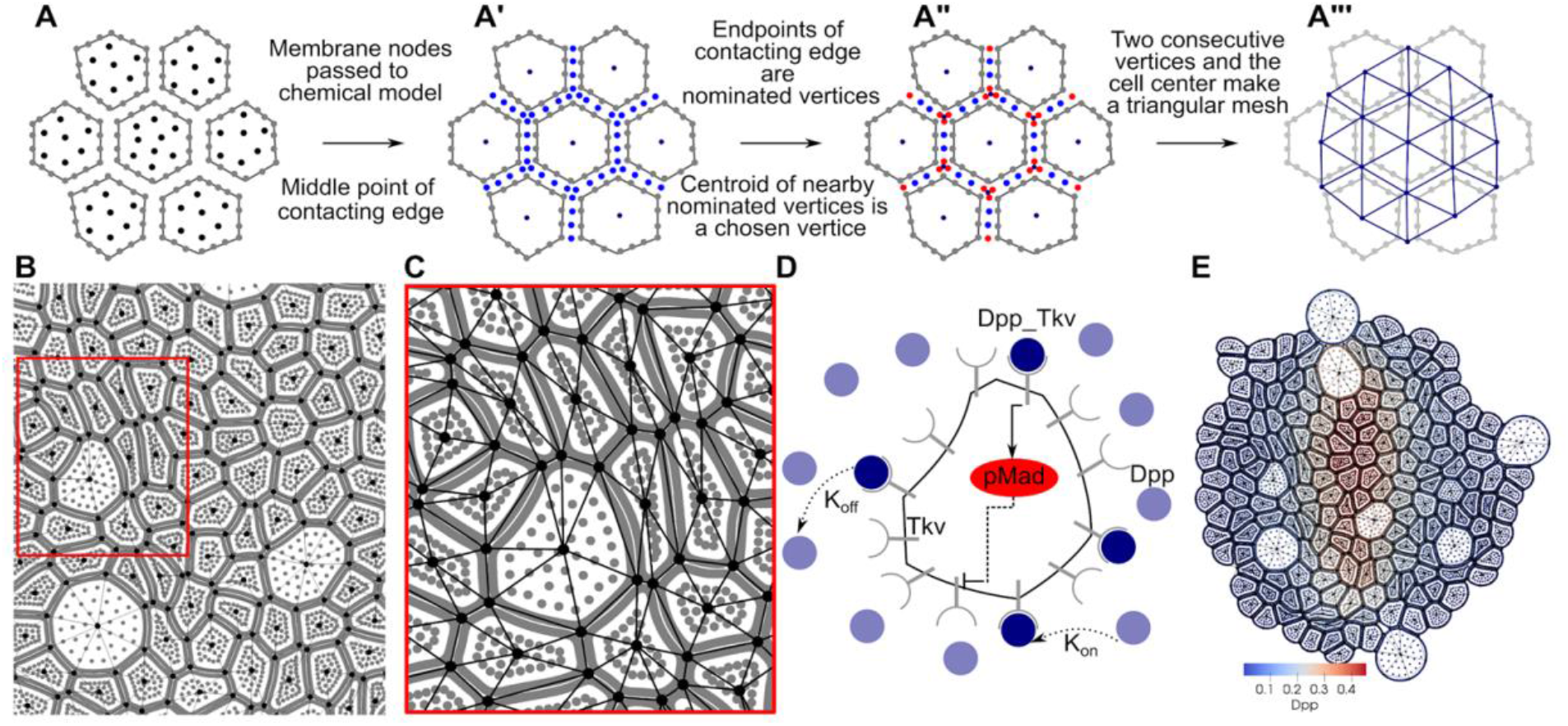
Spatial coupling of chemical and mechanical submodels. A) Nodes imported from the mechanical submodel. Black nodes represent cytoplasm and gray nodes represent cell membrane A’) Identification of common edges shared by neighboring cells. Blue dots are obtained as middle points of membrane nodes from neighboring cells. (A”) Identification of junction points among neighboring cells, denoted by red nodes. A’”) Triangulars obtained by connecting cell centers and junction points through common edges. B) Triangular mesh over a subset of the tissue. C) A zoom-in view of the resulting mesh. D) A schematic diagram of chemical signaling network within a single cell of Drosophila wing disc. E) Discretized tissue with Dpp gradient, denoted by blue-red color, obtained in the chemical-mechanical model.

#### Temporal Coupling of Mechanical and Chemical Submodels

The mechanical and chemical submodels were also coupled in time. Cell growth and division are usually initiated and regulated by some chemical signals. Moreover, the dynamics of chemical signals undergo a much faster time scale compared with mechanical changes. Therefore, a quasi-steady state of chemical signaling distribution was computed over the dynamic mesh capturing cell and tissue deformation, and transmitted into the mechanical submodel at some frequency, in order to guarantee consistent information exchanged. Such frequency had to be chosen appropriately, since too large frequency led to redundant computation and unnecessary computational cost, whereas too small frequency led to inconsistent chemical signals used in the mechanical submodel. In our model, change of the chemical signaling distribution depended on the deformation of individual cells. Therefore, To couple two submodels in time, we estimated the average time that one cell takes to enter the mitotic phase and divide. It was then used to estimate the frequency to update the quasi-steady state of chemical signaling distribution over the mesh constructed based on the most recent tissue configuration. More details were provided in Supplementary Information.

This multiscale chemical-mechanical model coupled in both time and space could be applied to simulate tissue development and investigate mechanisms underlying growth regulation. Next, we made specifications for the coupled model and applied it to study the development of *Drosophila* wing disc tissue.

#### Model specification for *Drosophila* wing disc

Dpp morphogen is the primary signal controlling cell growth and tissue development in *Drosophila* wing disc (Burke and Basler, 1996; Fried and Iber, 2014; Gibson and Perrimon, 2005; Harmansa et al., 2015; Matsuda and Affolter, 2017; Shen and Dahmann, 2005; Teleman and Cohen, 2000; Wartlick et al., 2011; Widmann and Dahmann, 2009; Zhou et al., 2012). In individual cells, Dpp binds with its receptors, Thickvein (Tkv), on the cell membrane to form the complex phosphorylates MAD (pMAD) as a downstream signal (Fig. 2D). Experimental data also suggest pMAD represses the production of Tkv as a negative feedback regulation (Zhu et al., 2020), leading to a lower synthesis of Tkv near the Dpp source region.

In the coupled model, dynamics of the morphogen and the intracellular signaling network were modeled by a system of reaction-diffusion equations following the structure of Eqn. 10–13 as below:

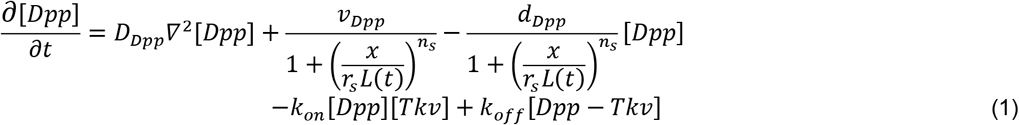

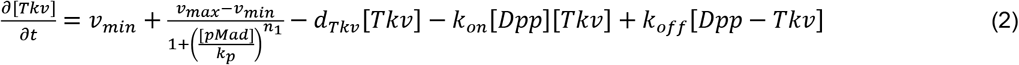

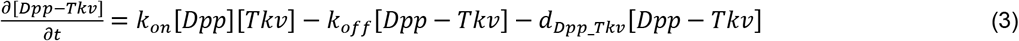

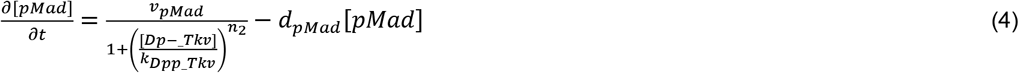

where *d*_*_’s represented degradation rates and *v*_*_’s represented production rates. *v_min_* and *v_max_* were the minimum and maximum production rates of Tkv receptors. The production of Dpp was modeled as a Hill function depending on the distance to the Dpp production region located at the AP boundary. In particular, the tissue width was denoted by *L*(*t*) and the width of the Dpp production region was denoted by *r_s_L*(*t*), where *r_s_* was a constant calibrated from experimental data (Zhu et al., 2020). The activation of intracellular signal pMad by the binding complex Dpp-Tkv was also modeled by a Hill function, so was the negative feedback regulation of pMad on Tkv.

For the cell division rule, we applied a hypothesis proposed in Wartlick et al., (2011) such that cells divide when the level of Dpp signal is increased by 50% compared with that at the beginning of each cell cycle, i.e.,

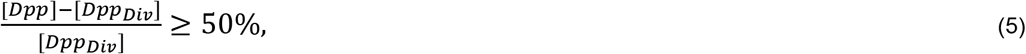

where [*Dpp*] represented the Dpp concentration at the current time and [*Dpp_Div_*] was the concentration at the beginning of one cell cycle. This hypothesis, also known as the temporal model, assumed that cells had a memory to keep track of Dpp level throughout the cell cycle and divided once the relative change of Dpp was sufficiently large. Based on this cell division rule, it was assumed in the coupled model that cells had a constant growth rate during the interphase (SI table 3), and they progressed into the mitotic phase if Eqn. 5 was satisfied (Fig. 5A).

**Figure 3.**
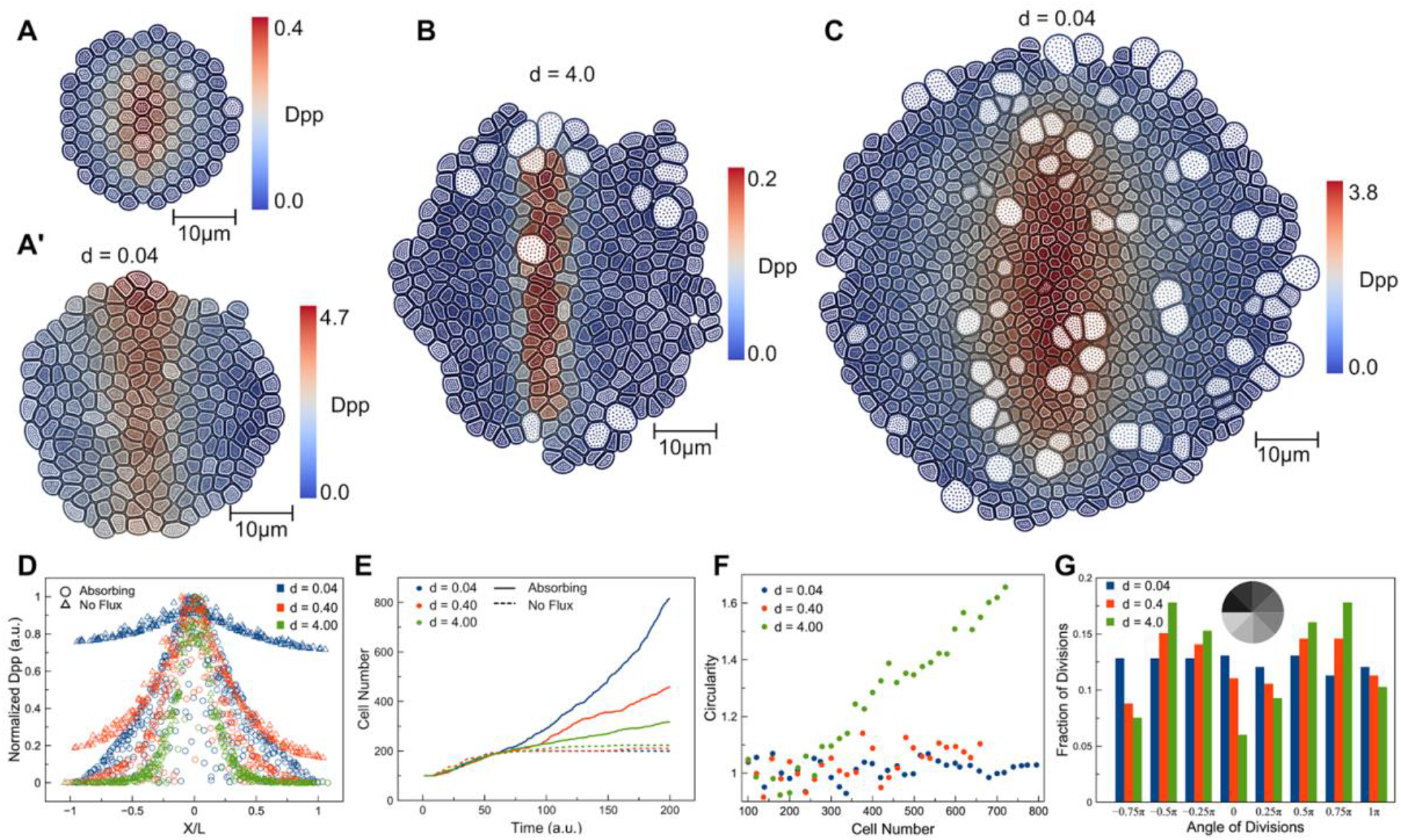
Simulation results of the coupled model with the simplified chemical submodel. A) Initial configuration of the tissue in simulations. Final configuration of the tissue at *t* = *200* with A’) no flux boundary condition and *d* = 0.04, B) absorbing boundary condition with large (*d* = *4.0*) or C) small (*d* = *0.04*) degradation rate of Dpp. D) Normalized Dpp profile at *t* = *200* with respect to the relative cell position in the tissue. E) Cell numbers with respect to time at different degradation rates and different boundary conditions. F) Tissue circularity with respect to the cell number at different degradation rates. Circularity was defined as the ratio of tissue height over tissue width. G) Distribution of the angular position of dividing cells with respect to tissue center for different degradation rates when there are *500* cells in the tissue.

**Figure 4.**
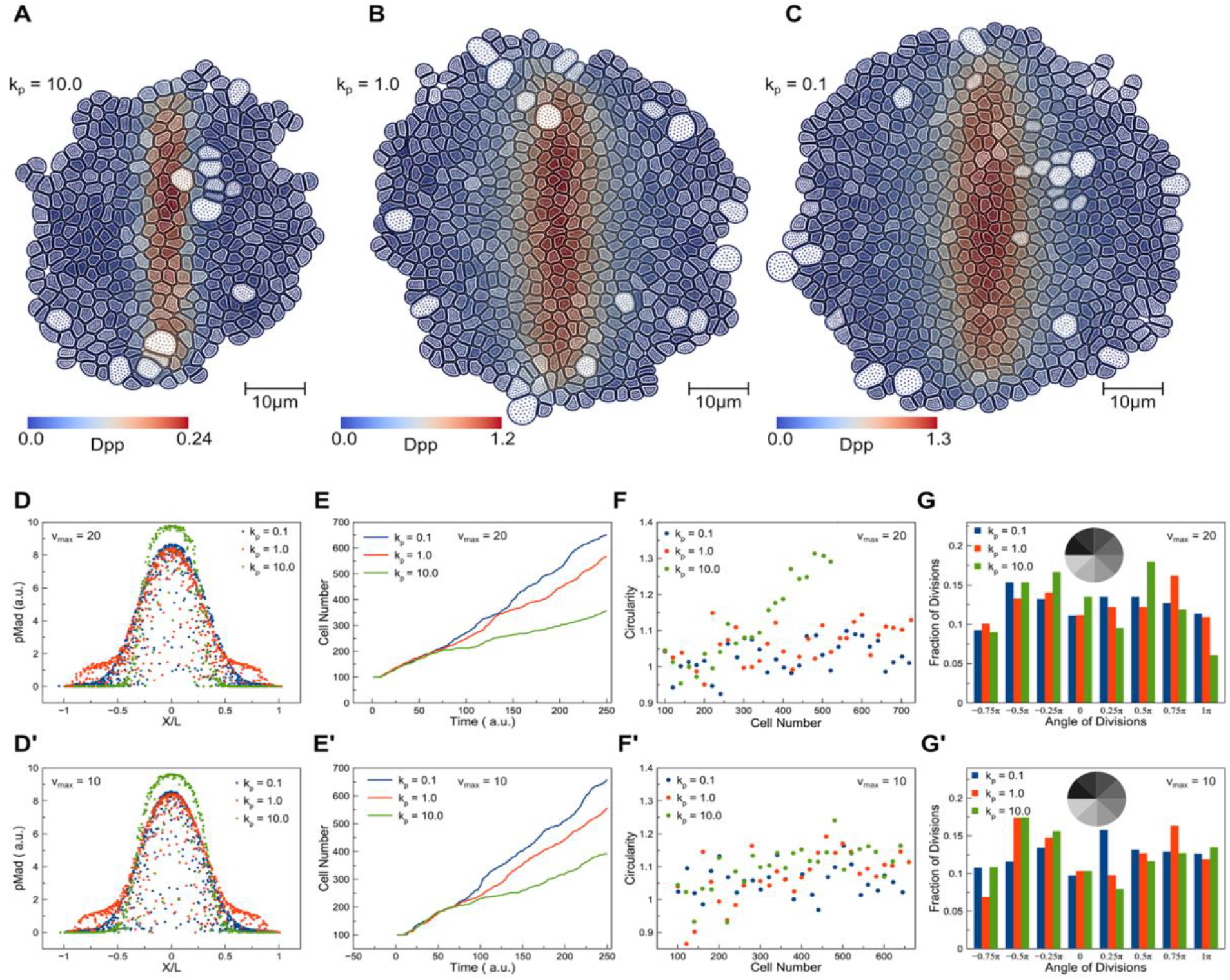
Simulation results with the advanced chemical submodel. Tissue configuration at *t* = *250* for *v_max_ = 20* with A) *k_p_* = *10.0* B) *k_p_* = *1.0* and C) *k_p_* = *0.1*. pMad profile at *t* = *250* with respect to the relative cell position in the tissue with different levels of feedback regulation and D) *v_max_* = *20* and D’) *v_max_* = *10*. Cell numbers at different levels of feedback regulation over time for E) *v_max_* = *20* and E’) *v_max_* = *10*. Tissue circularity with respect to cell number for different levels of feedback regulation for F) *v_max_* = *20* and F’) *v_max_* = *10*. Distributions of the angular position of dividing cells with respect to the tissue center for different levels of feedback regulation and G) *v_max_* = *20* and G’) *v_max_* = *10* when there are *500* cells in each simulation.

**Figure 5.**
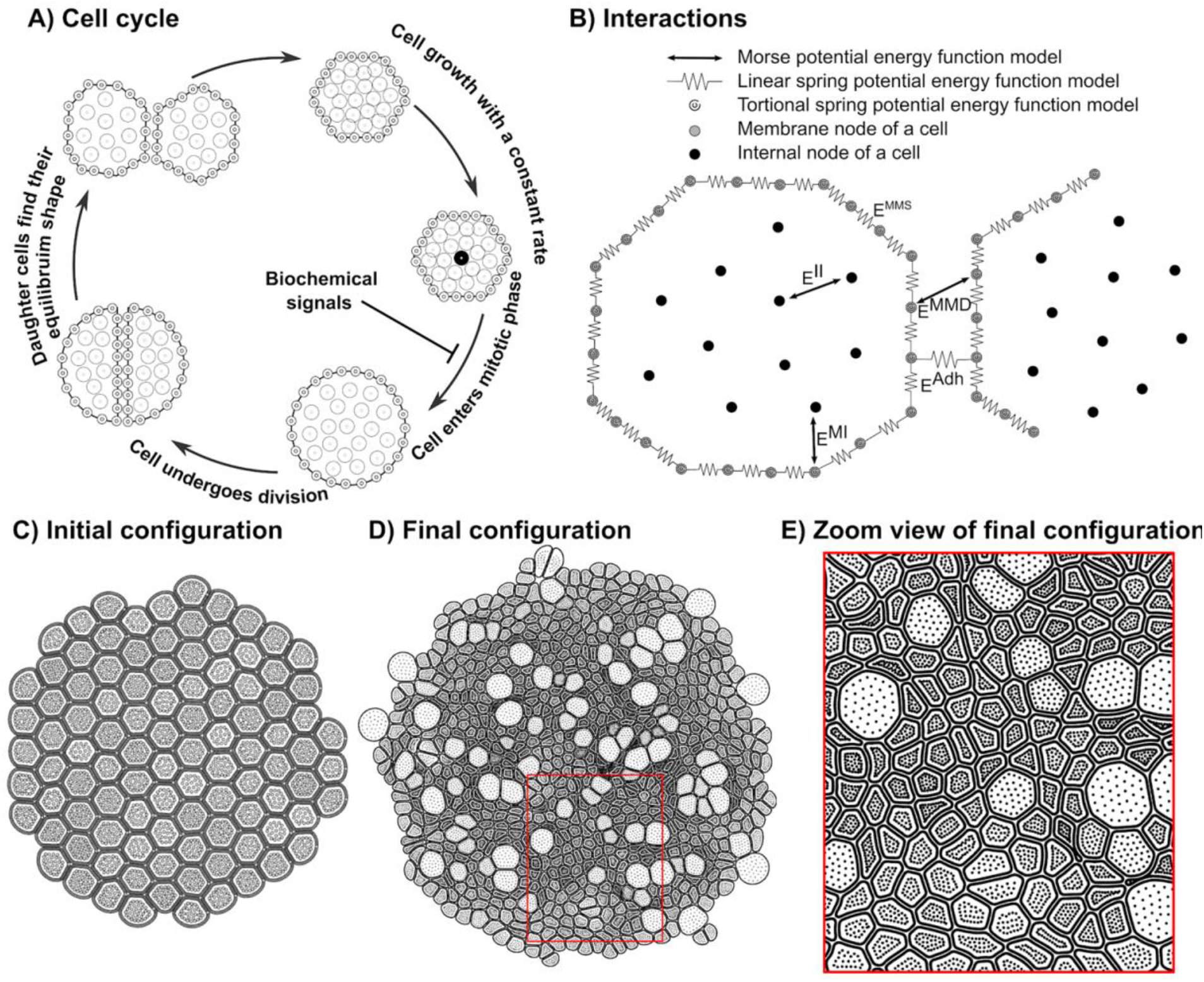
Diagram of the underlying physical basis of the mechanical sub-model. A) Life cycle of a cell in the mechanical submodel. B) Mechanical forces between different nodes in the subcellular element model. C) Initial tissue configuration in a simulation with no growth regulations. D) Final tissue configuration from the simulation in (C). E) Zoom-in view of the final configuration.

### Morphogen absorbance at the tissue boundary and large decay length prolong tissue growth at a fast and spatially homogeneous rate

Dpp is generated along the midline of the wing disc and diffuses bilaterally into the neighboring tissue. Therefore, it forms an exponential shape gradient along the AP axis (Fig. 1B). To characterize the Dpp gradient, it is common to use a quantity called decay length (*λ*), which measures the distance from the source region to the location where the Dpp level is reduced to *e^-1^* ≈ *37%* of its maximum (see section S5 for more details) (Zhu et al., 2020). A greater decay length represents a further spread of the exponential morphogen gradient. Interestingly, experimental data revealed that ubiquitous expression of Tkv led to a smaller decay length of the Dpp, followed by a slower growth and smaller tissue size (Zhu et al., 2020). This observation suggested the spatial distribution of morphogen gradient might play an important role in regulating tissue growth.

To understand how the distribution of the Dpp gradient affected tissue growth, we first considered a simplified chemical submodel by ignoring intracellular processes and downstream signals as below:

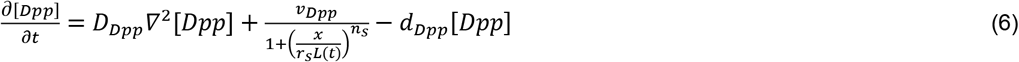

A reaction-diffusion equation with a specified source term along the midline (second term on the right-hand side of Eqn. 6) and a degradation term with a constant rate (the third term on the right-hand side of Eqn. 6) was adopted to capture a biologically relevant Dpp gradient. Moreover, the decay length of Dpp in this simplified model could be analytically estimated as 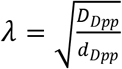 (See section S5 for details), depending on the diffusion rate and degradation rate. A higher diffusion rate or lower degradation rate allowed diffusive molecules to travel further, giving rise to a larger decay length. Both diffusion and degradation rates were calibrated to achieve a similar decay length observed in experiments (Zhu et al., 2020). We then coupled this simplified chemical submodel with the mechanical submodel under the specific cell division rule to simulate tissue growth. All simulations started with 100 cells as the initial condition (Fig. 3A), and the final shapes of simulated tissue development were shown in Fig. 3A’, B-C. Moreover, to understand how the decay length affected tissue growth, we perturbed the degradation rate of Dpp concentration in Eqn. 6, which changed the underlying decay length under different boundary conditions.

First, we considered the scenario that free Dpp molecules couldn’t escape at the boundary of the wing disc pouch and were always kept within the tissue. This was modeled by no flux boundary condition associated with cells located at the tissue boundary. For tissue growth with no flux boundary condition, smaller degradation gave rise to a flatter Dpp gradient (red and blue triangles in Fig. 3D). The Dpp concentration was saturated at high levels in individual cells and it didn’t increase sufficiently to satisfy the cell division rule at the early stage of development. Therefore, most cells only experienced one cell cycle, and tissue growth stopped at the early stage, leading to small tissue sizes (red and blue dash line in Fig. 3E). With a larger degradation rate, the Dpp gradient became more exponential (green triangles in Fig. 3D) and the final tissue size obtained was slightly increased (green dash line in Fig. 3E). However, tissue growth was still terminated early and the overall tissue size was much smaller than that obtained in experiments. Therefore, by assuming Dpp molecules couldn’t escape at the boundary, tissue growth only occurred in a short time period at the early stage and small final sizes were always obtained for different decay lengths of Dpp.

Second, we considered the scenario that Dpp was degraded completely at the periphery zone of the tissue (see section S3 for more information), which was modeled by absorbing boundary condition for cells at the boundary. For tissue growth with an absorbing boundary condition, the Dpp gradient changed from a linear shape to an exponential shape as the tissue size increased (circles in Fig. 3D). Furthermore, by assuming absorbing boundary conditions, the tissue growth was able to reach a size greater than the decay length of Dpp gradient (solid lines in Fig. 3E). We also observed that with a larger degradation rate, the morphogen gradient became exponential at a smaller tissue size, and the growth was maintained in a shorter period of time, giving rise to a smaller tissue size (solid green line in Fig. 3E). With a smaller degradation rate, the morphogen gradient became exponential at a larger tissue size (Fig. 3D), and the growth was maintained for a much longer time (Fig. 3E). These results were consistent with the experimental observation that a larger decay length of the Dpp gave rise to larger tissue sizes (Zhu et al., 2020).

In addition to limited growth, it was also noticed that with a higher degradation rate, the overall shape of the growing tissue, which was symmetric initially (Fig. 3A), became asymmetric, and the boundary became less smooth under the absorbing boundary condition (Fig. 3B). To look into that, we tracked the spatial locations of all dividing cells and visualized the distribution by dividing the tissue into eight sectors of equal size (Fig. 3G). It was observed that a higher degradation rate led to more dividing cells near the production region of Dpp, hence faster tissue growth along the AP boundary. As a result, the height of the tissue grew faster than the width, yielding an asymmetric shape (Fig. 3F). In contrast, a smaller degradation rate gave rise to more spatially homogeneous cell division (Fig. 3G) and a more symmetric overall tissue shape (Fig. 3F). Indeed, spatially homogeneous growth rate was also observed in experiments of *Drosophila* wing disc (Aegerter-Wilmsen et al., 2007; Bittig et al., 2009; Hufnagel et al., 2007; Milán et al., 1996; Vollmer and Iber, 2016; Wartlick et al., 2011), suggesting larger decay length of Dpp might be beneficial to achieve homogeneous growth in a wild-type wing disc tissue.

Overall, simulation results suggested that the decay length of the morphogen played an essential role in maintaining tissue growth and determining the final shape. Under the specific cell division rule, Dpp distribution with a larger decay length helped a tissue grow longer, faster, and in a more spatially homogeneous manner, which closely resembles the shape of the wild-type wing disc pouch observed in experiments. Absorbing boundary conditions with a lower degradation rate allowed Dpp molecules to travel further to establish a gradient with a larger decay length. In fact, it was observed that due to the hinge region around the wing disc pouch, Dpp level dropped to almost zero at the tissue boundary (Zhu et al., 2020), suggesting the absorbing boundary condition was more biologically relevant. Therefore, the absorbing boundary condition could also facilitate tissue growth by reshaping the morphogen gradient in a wild-type wing disc.

### Negative feedback on synthesis of receptors promotes the tissue growth through increasing morphogen decay length

It was well studied that the transduction of Dpp signals into cells in *Drosophila* wing disc relies on a receptor kinase Tkv. Removal of Tkv had a similar effect as the *Dpp* mutant. Recently, it was also observed that the intracellular downstream signal pMad downregulates the production of Tkv as a negative feedback regulation, which may reshape the morphogen gradient to some extent (Zhu et al., 2020). We applied our coupled model with absorbing boundary condition to study the effects of this feedback regulation on tissue growth.

Notice that in the chemical submodel (Eqns. 1–4), the negative feedback regulation of pMad on Tkv was modeled using a Hill function, as shown in Eqn. 2 and illustrated in Figure 2D. In particular, the parameter *k_p_* denoted the effective level of pMad involved in negative feedback regulation. Therefore, in simulations, we perturbed *k_p_* to give rise to different levels of the pMad feedback regulation. Higher *k_p_* values gave rise to weaker negative regulation in smaller regions, while lower *k_p_* values represented stronger negative regulation in larger domains. The cell division rule involved in the coupled model depended on pMad consequently.

Simulations were run for low (*k_p_* = 10), medium (*k_p_* = 1), and high (*k_p_* = 0.1) strength of negative feedback, as well as different values of maximal Tkv receptor production rate (*v_max_*) (Fig. 4). The simulation results showed that, with stronger strength of negative feedback of pMad (lower values of *k_p_*), the tissue grew faster (Fig. 4E, E’) and the overall shape of the tissue was more circular (Fig. 4F, F’). Moreover, the spatial distribution of dividing cells was more homogeneous (Fig. 4G,G’). However, it was also observed that simulation results for *k_p_* = 0.1 and *1* were similar to each other. This was because the production rate of Tkv became close to the minimum almost everywhere within the tissue for sufficiently small *k_p_*. Hence, the pMad gradient remained more or less the same for sufficiently small *k_p_*. By comparing the results generated using different values of the maximal receptor production rate (*v_max_* = 10 v.s. *20*), it was observed that the effect of the feedback regulation strength became more significant when *v_max_* was larger, i.e., the difference in the circularity of tissue shape due to different strength of the negative feedback regulation became more visible (Fig. 4F, F’).

In fact, stronger negative feedback regulation of pMad on Tkv receptors allowed Dpp molecules to diffuse into a larger area by reducing the binding occurrence on cell membrane, and therefore it gave rise to a pMad gradient with a larger decay length (Fig. 4D, D’). Based on the cell division rule used in this study, which depends on temporal changes of Dpp signal, a more widely spreading morphogen gradient helped tissue growth be maintained longer and cell numbers increase linearly at a faster rate, leading to a larger tissue size and more symmetric shape. This was also consistent with the results generated by the simplified chemical submodel (Eqn. 6), in which decreasing the degradation rate led to a larger tissue size and more spatially homogeneous cell division.

## Conclusions and Discussion

In this paper, we developed a multiscale coupled chemical-mechanical model to study growth regulation involved in tissue development, and applied it to study the development of *Drosophila* imaginal wing disc at the larva stage. The mechanical submodel described the shape change of individual cells and cell-to-cell physical interactions. It was coupled with a chemical submodel through an adaptive mesh generated over the growing tissue domain. This chemical signaling submodel described the dynamics of the morphogen gradient and associated downstream signals inside individual cells, which directed cell growth and division in the mechanical submodel. A specific cell division rule depending on the morphogen concentration sensed by individual cells was applied to understand how the decay length of the morphogen gradient affected tissue growth. By applying different boundary conditions in the chemical submodel, we found that tissue growth could be maintained longer under absorbing boundary conditions. This indicates that the significant reduction of morphogen at the hinge region surrounding a wing disc tissue could better promote tissue growth, compared with the case of the hinge region being a simple obstacle and preventing morphogen spread. By varying the decay length of the morphogen gradient, it was observed that the tissue grew faster with a greater decay length. Moreover, cell division became more spatially homogeneous, giving rise to a more symmetric tissue, consistent with experimental observations. We also found that the feedback regulation of pMad, a downstream signal of the morphogen, on the receptors indeed increased the decay length and therefore facilitated tissue growth. Overall, these results suggested the decay length of morphogen gradient could be critical in the hypothesized temporal model of growth regulation in the wing disc.

In this study, we applied a specific cell division rule based on the temporal changes of morphogen which was proposed in Wartlick et al., (2011). Our chemical-mechanical coupled model was developed as a general framework to study growth regulation of epithelial tissues, hence, it can be used to investigate other hypotheses on growth regulation. For example, it has been shown that cell mechanics contribute to growth control through a feedback loop in the wing disc (Shraiman, 2005; Hufnagel et al., 2007; Aegerter-Wilmsen et al., 2007), which might help to achieve a more uniform growth rate in the presence of an exponential morphogen gradient. In addition, it has been found that cell mechanics could regulate growth in a feedback loop by altering the Hippo pathway specifically (Pan et al., 2016). The coupled model is also extensible to study the cell growth rate as a function of both cell mechanical properties obtained from the mechanical submodel and chemical signals.

Since the mechanical and chemical submodels were coupled at the subcellular level in this work, it could potentially include a more complex interaction between mechanical components and chemical signals. For example, it was suggested that some signaling pathways could be affected by cell mechanical properties including shear stress and tension sensed at adherens junctions (Dupont, 2016; Irvine and Shraiman, 2017; Sun and Irvine, 2016). Meanwhile, signaling molecules could rearrange structural components within individual cells and direct new materials to the cell membrane to modify the mechanical properties (Widmann and Dahmann, 2009). These interactions between chemical and mechanical components can be incorporated into the coupled model upon availability of more detailed experimental quantification and tested for their roles in cell growth regulation and tissue development.

## Materials and Methods

### Mechanical submodel

For the mechanical submodel, we follow a similar approach as the Epi-scale model (Nematbakhsh et al., 2017). Epi-scale model is a multiscale subcellular element computational platform that simulates the growth of epithelial monolayers with detailed cell mechanics. Individual cells are represented as collections of two types of interacting subcellular nodes: internal nodes and membrane nodes. Internal nodes account for the cytoplasm and the membrane nodes represent both the plasma membrane and associated contractile actomyosin cortex. Interactions between internal and membrane nodes are modeled by using potential energy functions, as shown in Fig. 5B (Christley et al., 2010; Newman, 2005). Combined interactions between pairs of internal nodes (*E^II^*) represent the cytoplasmic pressure of a cell. Combined interactions between internal and membrane nodes of the same cell (*E^MI^*) represent the pressure from the cytoplasm to the membrane. Interactions between membrane nodes of the same cell (*E^MMS^*) represent cortical stiffness. Cell-cell adhesion (*E^Adh^*) is modeled by combining pairwise interactions between membrane nodes of two neighboring cells. *E^MMD^* is a repulsive force between membrane nodes of neighboring cells and prevents membranes from overlapping. Springs and Morse energy functions are utilized to model all the interactions (Sandersius and Newman, 2008). The following equations of the motion describe movements of internal and membrane nodes, respectively:

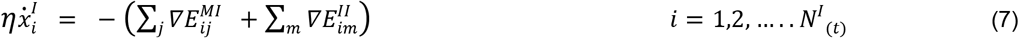

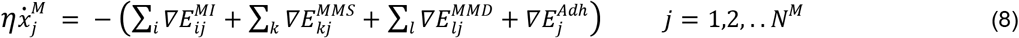

where *η* is the damping coefficient, 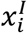 and 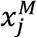 are positions of internal and membrane nodes. *m* is the index for any internal node. *k* is the index for any membrane node of the same cell interacting with the membrane node *j*. *l* is the index for any membrane node of a different cell interacting with the membrane node *j*. Cell growth is modeled by adding internal nodes meaning that *N^I^* increases based on cell proliferation rate. The individual cell cycle in the current model is shown in Fig. 5A. Initial and final configurations of the tissue in a simulation with a given growth rate and cell division rate are shown in Fig. 5C and Fig. 5D,E, respectively.

### Chemical submodel

In the chemical signaling model, we consider a chemical signal which regulates the growth rate and cell division. A morphogen, which is a signaling molecule governing the growth and patterning of tissue development, diffuses in the extracellular space to form a gradient at the tissue level. A reaction-diffusion equation is used to model the spatiotemporal morphogen dynamics as below:

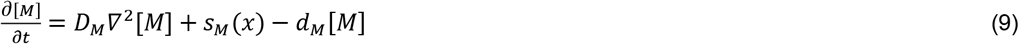

where [*M*] denotes the concentration of the morphogen molecules, *D_M_* is the diffusion coefficient of morphogen molecules. *d_M_* is the degradation rate of morphogen molecules. The production rate of morphogen molecules, varying spatially, is denoted by *S_M_*(*x*). *D_M_* and *d_M_* together determine how far the molecules can reach in the steady state (See section S5 for more information). The local morphogen concentration is sensed by individual cells through binding with receptors on the cell membrane to activate the intracellular signaling network.

To model the intracellular signaling network, we consider a receptor *R*, the complex after binding *MR*, and a downstream signal *S*. More components with more complex regulations can be modeled similarly. Together with the diffusive morphogen, it gives rise to a chemical signaling network at both cell and tissue levels, formulated as Eqn. 10–13 below. More specifically, the binding of the morphogen molecules and receptors is reversible, so both binding and unbinding kinetics are included with *k_on_* denoting the binding rate and *k_off_* characterizing the unbinding rate. A standard Hill function is applied to model the activation of downstream signal S by the complex *MR*. Maximal signal production rate and concentration at which the production is half of the maximum are denoted by *v_s_* and *k_MR_*, respectively. It is assumed that S regulates the synthesis of the receptor as a feedback regulation, which is also modeled as a Hill function, to accommodate the feedback regulation present in the *Drosophila* wing disc. The minimum and maximum of receptor production rate are *v_R,min_* and *v_R,max_*. The concentration producing half occupation is represented by *k_s_*. Notice that only *M* can diffuse within the tissue, and all other components are restricted within the cell without diffusion.

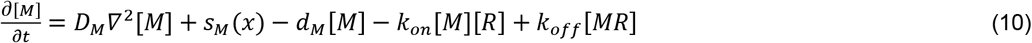

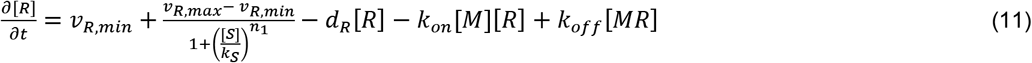

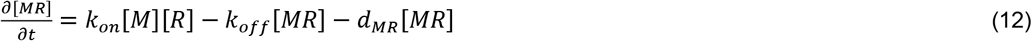

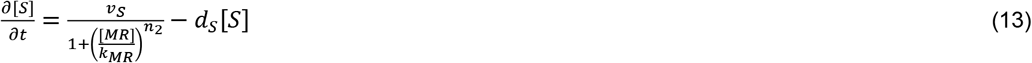

### Dynamic mesh generator to couple mechanical and chemical submodels

To generate a computational mesh for the chemical signaling submodel, we first identify neighbors of individual cells based on the distance between membrane nodes of every two cells (See Fig. 2A and section S2 for more information). In particular, one cell is considered to be a neighbor of the other if the shortest distance between their membrane nodes is less than some threshold. This threshold is chosen based on the distance between neighboring cells obtained in the equilibrium in the simulation (See section S2 for more details). The same threshold is also used to determine a common edge between neighboring cells, i.e., membrane nodes from neighboring cells with a distance less than the threshold are selected to calculate a common edge. Then middle points are calculated for each pair of those selected nodes, which give rise to a common edge between these two neighboring cells (Fig. 2A’). The endpoints of each shared edge are used to determine vertices of the triangular mesh. It is possible that multiple cells neighboring to each other give rise to a junction. Therefore we consider all common edges associated with the same junction point and calculate the centroid of their endpoints near the junction as a vertex in the triangular mesh (red dots in Fig. 2A’’). We go over all junctions and calculate corresponding vertices throughout the tissue. Next, the center of each cell is calculated as the centroid of all its membrane nodes, and it is connected to vertices obtained at junctions (Fig. 2A’”). By doing that, each cell is discretized by a triangular mesh that shares a common edge with its neighboring cells, and triangles in all cells give rise to a mesh covering the entire tissue (Fig. 2B,C). Notice that boundary cells usually lack neighbors along one or more sides, therefore their discretization will be treated separately (See section S2.1 for more information). Nodes from cell membranes that act as the tissue boundary in those cells are selected as vertices to satisfy some minimal distance between successive ones. They are denoted by boundary vertices and connected with the corresponding cell center to give rise to the triangular mesh inside boundary cells. A mesh quality check is implemented to guarantee that no highly skewed triangles are generated for convergence and accuracy of the computation over the mesh. Adjustment is conducted by merging or splitting triangles if triangles are found to be too skewed in the quality check (See section S2 for more information). Such a mesh generator provides triangular meshes in individual cells, as well as a global mesh over the whole tissue. Moreover, the triangular mesh is updated at some frequency to accommodate the cell deformation and tissue growth obtained in the mechanical submodel.

## Acknowledgment

This work used the Extreme Science and Engineering Discovery Environment (XSEDE) PSC Bridges GPU (Bridges GPU) at the service-provider through allocation DMS200024. XSEDE is supported by National Science Foundation grant number ACI-1548562. Simulations were also partially performed using the resources of the HPCC at University of California, Riverside. We thank Nael Abu Ghazaleh and Hodjat Asghari Esfeden for providing additional computational resources. We thank Jeremiah Zartman for suggestions and discussion.

## Data availability

https://github.com/arame002/EpiScale_Signal.git

## Author contributions

Conceptualization: W.C., A.N.; Methodology: W.C., A.N., M.A., R.Z.; Formal analysis: A.R., A.N., W.C.; Investigation: A.R., A.N., W.C.; Software: A.R., S.B., W.C., A.N.; Visualization: A.R., W.C., A.N.; Writing: W.C., A.N., M.A., R.Z.; Supervision: W.C., A.N.; Project administration: W.C., A.N.; Funding acquisition: W.C., A.N., M.A., R.Z.

